# Iron regulatory proteins 1 and 2 have opposing roles in regulating inflammation in bacterial orchitis

**DOI:** 10.1101/2023.08.16.553579

**Authors:** N. Ghatpande, A. Harrer, B. Azouly, N. Guttmann-Raviv, S. Bhushan, A. Meinhardt, E.G. Meyron-Holtz

**Affiliations:** Faculty of Biotechnology and Food Engineering, Technion-Israel Institute of Technology, Technion City, Haifa, Israel; Institute of Anatomy and Cell Biology, Unit of Reproductive Biology, Justus-Liebig-University of Giessen, Giessen, Germany

**Author notes:** equal last and corresponding authorship. equal first authorship (N. Ghatpande, A. Harrer).

## Abstract

Acute bacterial orchitis (AO) is a prevalent cause of intra-scrotal inflammation, often resulting in sub-or infertility. A frequent cause eliciting AO is uropathogenic *Escherichia coli* (UPEC), a gram negative pathovar, characterized by the expression of various iron acquisition systems to survive in a low-iron environment. On the host side, iron is tightly regulated by iron regulatory proteins (IRP) 1 and 2 and these factors have been reported to play a role in testicular and immune cell function, however, their precise role remains unclear. Here, we show in a mouse model of UPEC-induced orchitis that the absence of IRP1 results in reduced immune response and testicular damage. Compared to infected wild-type (WT)-mice, testis of UPEC-infected *Irp*1^-/-^ mice showed impaired ERK signalling. Conversely, IRP2 deletion led to a stronger inflammatory response. Notably, differences in immune cell infiltrations were observed among the different genotypes. In contrast to WT and *Irp*2^-/-^ mice, no increase in monocytes and neutrophils was detected in testis of *Irp*1^-/-^ mice upon UPEC-infection. Interestingly, in *Irp*1^-/-^ UPEC-infected testis, we observed an increase in a subpopulation of macrophages (F4/80+ CD206+) associated with anti-inflammatory and wound-healing activities compared to WT. These findings suggest that IRP1 deletion may protect against UPEC-induced inflammation by modulating ERK signalling and dampening the immune response.

## Introduction

Acute orchitis (AO) is a prevalent cause of intra-scrotal inflammation that results in approximately 600,000 medical visits per year in the United States alone and presents mostly as combined epididymo-orchitis (Rupp and Leslie, 2023). AO is predominantly attributed to uropathogenic *Escherichia coli* (UPEC) strains or sexually transmitted pathogens such as *Chlamydia trachomatis* (Nicholson et al., 2010; Pilatz et al., 2015). This condition can result in infertility in males, and studies have shown that patients infected with UPEC have lower sperm counts even many months after successful antibiotic treatment (Bhushan et al., 2009; Fraczek and Kurpisz, 2015; Pilatz et al., 2015; Schuppe et al., 2017).

Iron, an essential element for humans, plays an important role in spermatogenesis (Tvrda et al., 2015). Mammalian iron homoeostasis is tightly regulated at both the systemic and cellular levels. IRP1 and IRP2 (also known as ACO1 and IREB2, respectively) are crucial for maintaining cellular iron homoeostasis (Rouault, 2006; Wilkinson and Pantopoulos, 2014). This tight regulation ensures the balance of cellular iron levels (Muckenthaler et al., 2008). IRP2 deletion has been found to significantly alter iron metabolism, highlighting its critical role in cellular iron homoeostasis, whereas IRP1 deletion impairs iron metabolism only slightly and in a tissue-specific manner (Ghosh et al., 2013; LaVaute et al., 2001; Wilkinson and Pantopoulos, 2013). In addition to their role in cellular iron homoeostasis, recent studies have shown their roles in immune regulation as well. For example, Bonadonna et al. (2022) showed that IRPs ensure that neutrophils have the iron they need to fight infection, and their deficiency leads to impaired neutrophil function and increased susceptibility to bacterial infections (Bonadonna et al., 2022). Furthermore, ablation of IRP1 and IRP2 in macrophages leads to increased susceptibility to *Salmonella* infection, suggesting that these proteins are critical for limiting microbial iron acquisition and promoting host defence (Nairz et al., 2015). However, the precise role of IRPs in the immune response remains poorly understood.

Iron is also an essential nutrient for the growth and survival of most bacterial species. During bacterial infection the host typically limits systemic iron availability to pathogens as a part of the immune response (Haschka et al., 2021; Nairz and Weiss, 2020). However, certain bacterial species, such as UPEC, have evolved specialised mechanisms to acquire iron from the host, potentially contributing to their virulence (Dabral et al., 2022; Frick-Cheng et al., 2022; Robinson et al., 2018). Further studies have shown that limiting iron can reduce the bacterial burden (Bauckman et al., 2019). To circumvent the low iron availability, UPEC can chelate iron via siderophores from the host. Alternatively, UPEC can persist in cells within autophagosomes, where they can access free iron via ferritinophagy (Bauckman and Mysorekar, 2016). In addition, in bladder infection, UPEC induced activation of Toll like receptor 4 (TLR4), which is known to lead to the production of pro-inflammatory cytokines, including IL-6 and IL-1*β* (Schilling et al., 2001; Song et al., 2007). In contrary, in rat testis, pro-inflammatory cytokines like TNFα and Il-6 were not produced following UPEC infection (Bhushan et al., 2008). Furthermore, modified iron availability during bacterial infections influences the polarisation of macrophages into classically activated (M1) or alternatively activated (M2) macrophages pending upon location of infection (Behmoaras, 2021; Ni et al., 2022).

M2 macrophages, characterized e.g. by the high constitutive production of immunosuppressive cytokines such as IL-10, play a relevant role in normal testicular homoeostasis (Bhushan et al., 2015; Wang et al., 2017). On the other hand, in orchitis infiltrating monocytes and neutrophils are instrumental in causing the observed tissue damage (Bhushan et al., 2011; Klein et al., 2020; Wagenlehner and Naber, 2006; Wang et al., 2021). In this regard, it is completely unknown what role iron homoeostasis plays in the development of orchitis and in the magnitude of testicular immune response. Thus, in this study we aimed to investigate the role of iron homoeostasis on UPEC-mediated orchitis in mice with targeted deletions of IRP1 or IRP2.

## Results

### IRP1 deficiency protects the testis against UPEC-induced inflammation

To examine whether inflammatory effects of UPEC-infection were exacerbated or attenuated in *Irp*1^-/-^ and *Irp*2^-/-^ mice compared to wild type (WT), UPEC was injected into both *vasa deferentia* of mice close to the epididymides and effects were monitored after 7 days in testis (Fig. 1A). Testicular weight was similar in sham and UPEC-infected mice of all three genotypes (suppl. Fig. 1E). Yet, H&E-stained testicular sections revealed notable alterations in testicular morphology in WT and *Irp*2^-/-^ infected mice (Fig. 1B, green arrowheads). Surprisingly, *Irp*1^-/-^ mice exhibited predominantly normal spermatogenesis following UPEC-infection, even though bacterial loads in *Irp1*^*-/-*^ mice were comparable to those in WT and *Irp2-/-* mice (suppl. Fig. 1F, G).

**Figure 1:**
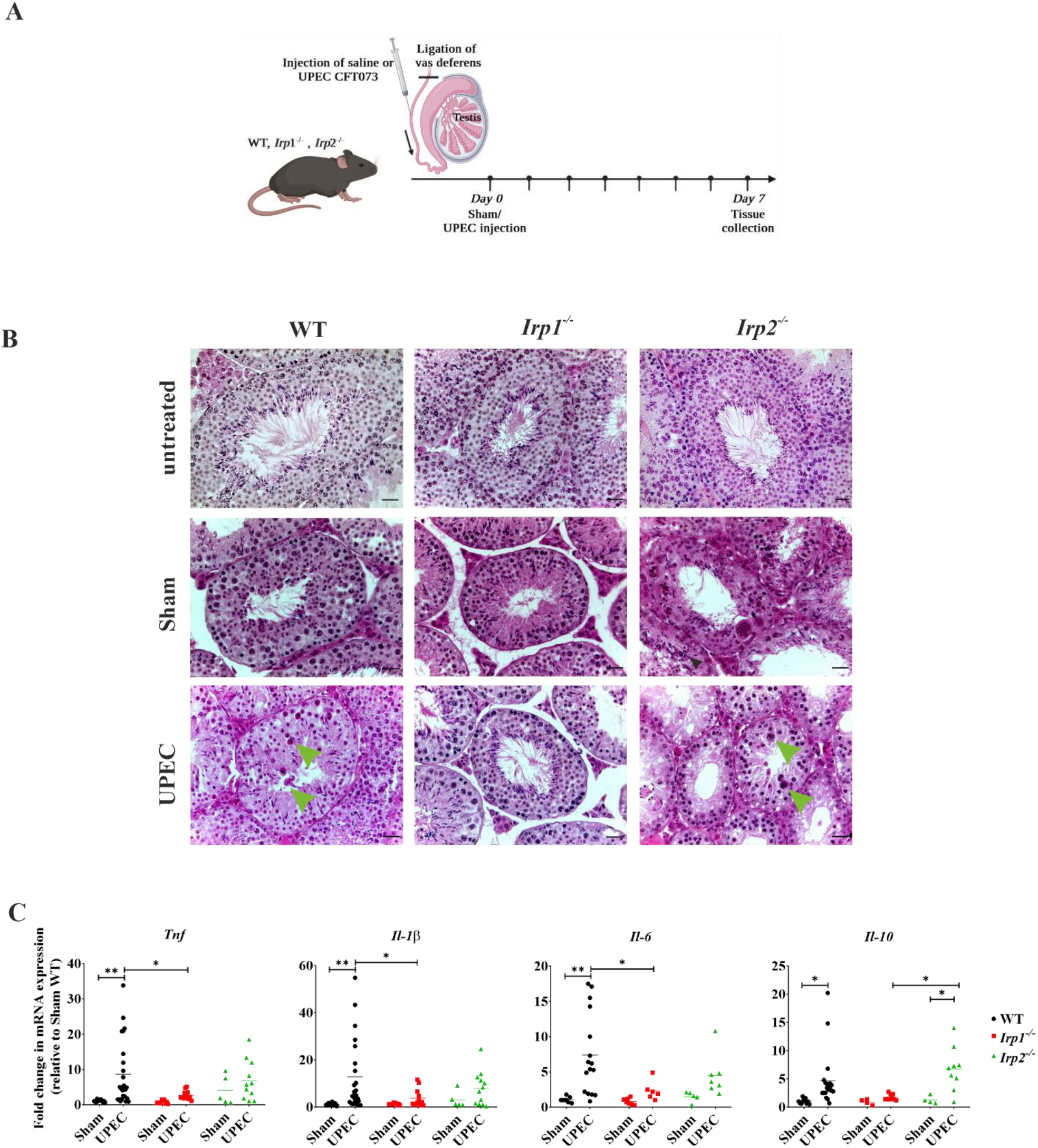
IRP1 deficiency protects the testis against UPEC-induced inflammation. (A) Experimental design illustrating the UPEC-induced orchitis mouse model (created in BioRender.com). WT, *Irp1*^*-/-*^ and *Irp2*^*-/-*^ mice were injected with saline or UPEC CFT073 via the vas deferens. Organs were collected seven days post-infection for further analysis by histology, flow cytometry and quantitative RT-PCR. (B) Histopathological analysis of UPEC-infected WT and *Irp*2^-/-^ testis revealed impairment of spermatogenesis including multinucleated cells (green arrowheads). Representative micrographs of hematoxylin and eosin stained testes are shown. Scale bar: 20 μm (N=5-7). (C) Quantitative RT-PCR analysis demonstrated altered expression levels of key pro-inflammatory (*Tnf, Il1-β, Il-6*) and anti-inflammatory (*Il*-*10*) cytokines in UPEC-infected testes. Relative mRNA levels were normalised to *Rplp*0 and further to sham WT. FC= Fold change. Statistical significance was determined using two-way ANOVA with Tukey multiple comparison (\**P*<0.05, ** *P*<0.01).

To gain further understanding of the observed decrease in testicular damage in UPEC-infected *Irp1*^*-/-*^ mice, we analysed possible alterations in immune response by quantifying the mRNA expression levels of key pro-inflammatory (*Tnf, Il-6*, and *Il-1β*) and anti-inflammatory (*Il-10*) cytokines using quantitative PCR (qPCR), where a significant increase in cytokine transcript levels in UPEC-infected WT and *Irp*2^-/-^ testes were noted compared to sham-infected testes (Figure 1C).

### Iron status remains unchanged after UPEC infection

As UPEC infection can mediate significant alterations of the iron status (Bauckman et al., 2019), we examined cellular iron status in the testis, seven days after UPEC infection and focused on proteins regulated by IRPs. No significant changes in the expression levels of the ferritin subunits (FtH and FtL) and transferrin receptor 1 (*Tfr1*) were observed in response to the infection across different genotypes on both the protein and mRNA level, respectively (Supp. Fig. 2A-D). Further analysis using immunofluorescence staining on WT mouse testis revealed that ferritin was predominantly localised within the interstitium, primarily in macrophages (supp. Fig. 2E).

### The ERK signalling pathway is impaired in *Irp1*^*-/-*^ testis

As the absence of an inflammatory response in *Irp*1^-/-^ mice could not be explained by differences in the iron status, we asked whether our observation could be explained by a modified activation of the proinflammatory MAPK and NF-kB pathways that are established to play a role in UPEC elicited testicular inflammation (Bhushan et al., 2008; Krachler et al., 2011; Wang et al., 2009). Levels of TLR4 and p-P65 (NF-kB pathway) remained unchanged among all genotypes. In contrast, levels of phosphorylated ERK (p-ERK) were significantly lower in response to UPEC infection in *Irp1*^*-/-*^ testis compared to WT organs, whereas p-P38 remained unchanged (Fig. 2A,B). This was confirmed in bone marrow derived macrophages (BMDM) stimulated with LPS where p-ERK levels were increased in WT and *Irp*2^-/-^ BMDM with only a moderate increase seen in BMDM from *Irp1*^*-/-*^ mice (Fig. 2C, D). Basal levels of p-ERK in BMDM of *Irp*2^-/-^ mice were significantly higher than in WT and *Irp*1^-/-^ mice. No significant differences in phosphorylation of P38 upon LPS treatment were observed across all genotypes (Fig. 2E).

**Figure 2:**
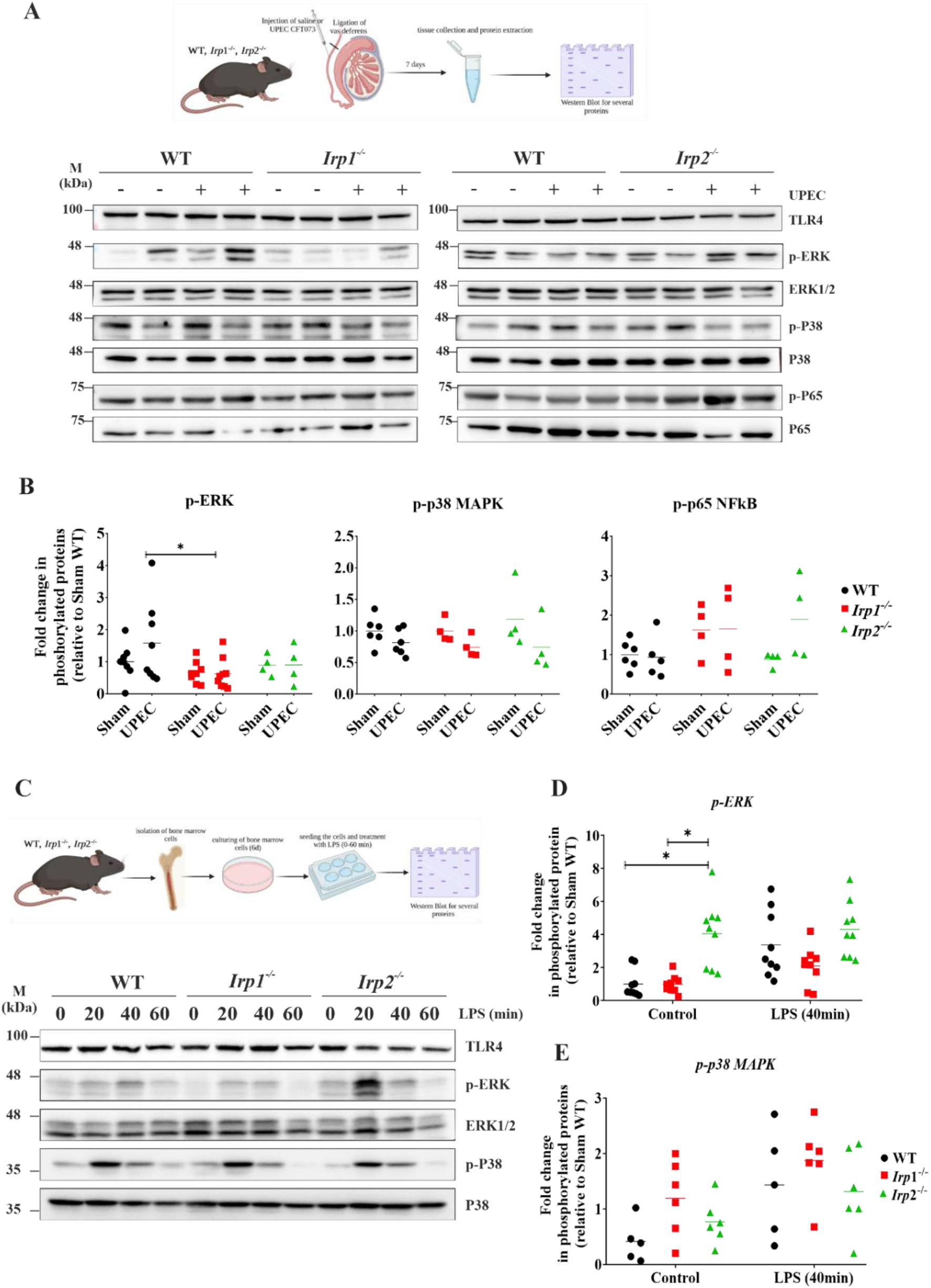
Impairment of the ERK signalling pathway in *Irp1*^*-/-*^ testis. (A-E) Protein expression changes of various signalling proteins were analysed by Western blot. (A) The experimental workflow is shown (created in BioRender.com). Levels of targeted proteins (TLR4, p-ERK, total ERK, p-P38, P38, p-P65 and P65) are shown. The band intensities of p-ERK, p-P38, and p-P65 (B) were quantified using ImageJ and normalised to the corresponding loading control. Statistical significance was determined using two-way ANOVA with Tukey multiple comparison (N=6; \**P*<0.05, ** *P*<0.01). (C) Bone marrow-derived macrophages (BMDM) from WT, *Irp1*^*-/-*^ and *Irp2*^*-/-*^ mice were isolated and treated with 200 ng/ml lipopolysaccharide (LPS) for different time points (20, 40 and 60 min) under 6% O2 conditions at 37°C. Extracted proteins from isolated BMDM were analysed by Western blot. Representative blots from three independent experiments are shown. (D, E) Quantitative analysis was performed as described above. Statistical significance was determined using two-way ANOVA with Tukey multiple comparison (\**P*<0.05, ** *P*<0.01).

### Infiltration of immune cells is lower in *Irp*1^-/-^ testis compared to WT and *Irp2*^*-/-*^ mice

We speculated that the reduced p-ERK signalling observed in infected *Irp*1^-/-^ mice could be accompanied with changes in leukocytic infiltration in the testis during UPEC infection (Talbot et al., 2011). Flow cytometry analysis showed no significant differences in total leukocytes (CD45^+^ cells) in naive mice among genotypes (suppl. Fig. 1A, B). Similarly, in the blood no significant differences in leukocytes were observed across genotypes (suppl. Fig. 1D). However, we found a slight reduction in the proportion of macrophages (F4/80^+^CD11b^+^) in total CD45^+^ leukocytes in naive-*Irp*2^-/-^ mice as compared to naive-WT and *Irp*1^-/-^ mice (suppl. Fig. 1C). Circulating macrophages (F4/80^+^CD11b^+^) in the blood did not exhibit such differences (suppl. Fig. 1D).

As seen previously during UPEC infection, we observed an increase in total (CD45^+^) leukocytes in WT testis and found a similar pattern in testis of *Irp*2^-/-^ mice (Fig. 3A). However, in UPEC-infected *Irp*1^-/-^ testis no changes in the percentage of CD45^+^ cells were seen compared to sham control (Fig. 3A).

**Figure 3:**
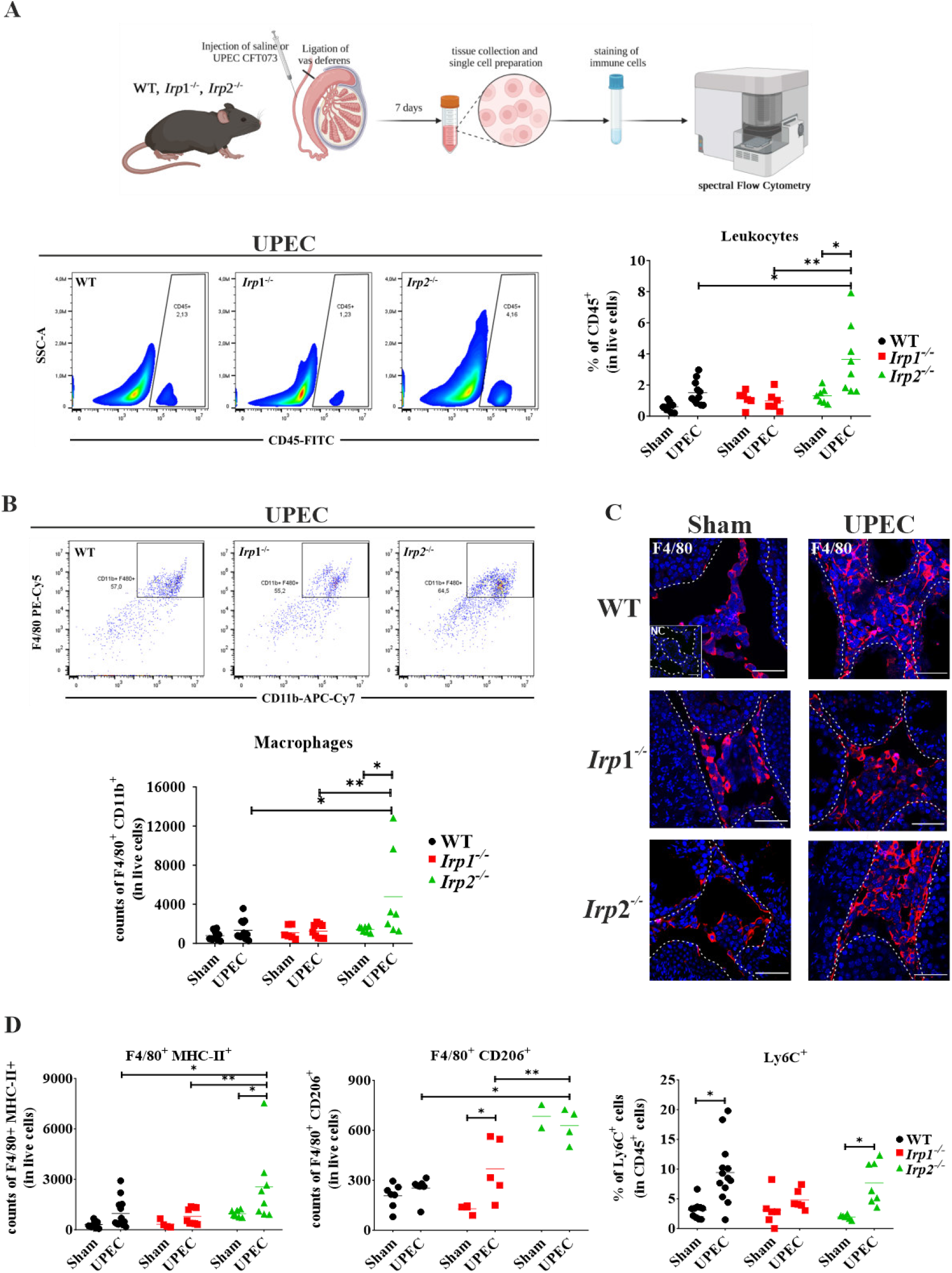
UPEC-induced leukocyte infiltration in the testis is lower in *Irp1*^*-/-*^ as compared to WT and *Irp2*^*-/-*^. (A) The scheme shows the experimental workflow for flow cytometry analysis (created in BioRender.com). Representative flow cytometry plots and corresponding bar graphs depict the total leukocyte population (CD45^+^, in 2×10^5^ live cells) after UPEC infection. (B, C) Detailed analysis of total macrophages (CD11b^+^ F4/80^+^), (D) subpopulations of macrophages (F4/80^+^MHC-II^+^; F4/80^+^CD206^+^) and monocytes (Ly6-C^+^) in testes of different mouse genotypes. Representative flow cytometry plots and corresponding graphs are shown. (C) F4/80 (red) immunofluorescence was shown to localise macrophages in the testes of all genotypes. Data were obtained from 6-9 mice per group. Scale bar represents 50 μm.

Further flow cytometry analysis revealed no increase in total macrophage numbers (F4/80+ CD11b+) in UPEC-infected *Irp1*^*-/-*^ testis, while in *Irp*2^*-/-*^ mice a strong increase was observed (Fig. 3B). Immunofluorescence analyses supported the view from flow cytometry about increased numbers of F4/80^+^ cells within the interstitium of the WT and *Irp*2^*-/-*^ testis after UPEC infection (Fig. 3C).

The MHC-II^+^ macrophage subpopulation showed a similar elevation in UPEC-infected WT and *Irp*2^-/-^ testis but not in *Irp*1^-/-^ testis (Fig. 3D). However, the subpopulation of CD206^+^ macrophages was elevated exclusively in UPEC-infected *Irp*1^-/-^ testis compared to sham *Irp*1^-/-^ testis (Fig. 3D). This is important as CD206^+^ macrophages play a crucial role in immunoregulation and in the resolution of tissue inflammation in the testis and other organs (Bhushan et al., 2020). Concomitant to macrophages, the monocytes (Ly6C^+^ cells) also showed a significant increase only in WT and *Irp*2^-/-^ testis but not in *Irp*1^-/-^ testis after UPEC infection (Fig. 3D). Neutrophil numbers (Ly6G^+^) in *Irp*2^-/-^ testis were also elevated post UPEC infection compared to WT and *Irp*1^-/-^ as demonstrated by flow cytometry and immunofluorescence analysis (Fig. 4A, B and C), whilst no differences in T cell (CD3^+^) numbers across genotypes were evident. B cell (CD19^+^) numbers decreased in WT-infected testis compared to the respective sham control (suppl. Fig. 3C and D).

**Figure 4:**
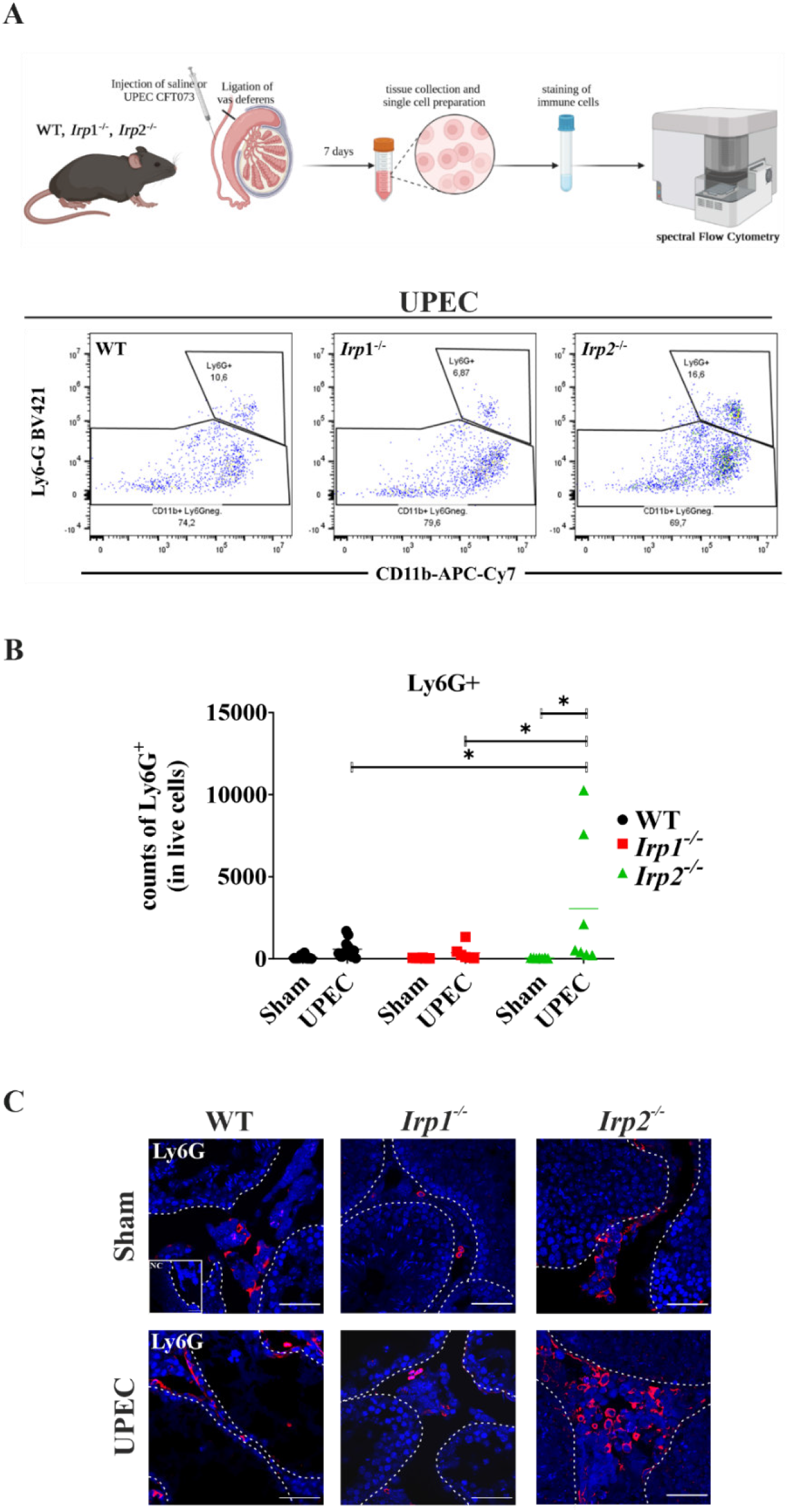
Increased neutrophil recruitment in infected *Irp2*^*-/-*^ testis compared to WT and *Irp1*^*-/-*^ mice. (A) The scheme shows the experimental workflow for flow cytometry analysis (created in BioRender.com). Representative flow cytometry plots and corresponding bar graph (B) demonstrate the differences in neutrophil counts across the different genotypes. Statistical significance was determined using two-way ANOVA with Tukey multiple comparison (\**P*<0.05, ** *P*<0.01). (C) Immunofluorescence staining of Ly6-G was performed to visualize the localisation of neutrophils in UPEC infected testes.

Further analysis on chemokine expression related to monocyte, macrophage (*Ccl2*), and neutrophil (*Cxcl2*) recruitment revealed lower expression of *Ccl2* and *Cxcl2* in testis from UPEC-infected *Irp1*^*-/-*^ mice compared to testis from UPEC-infected WT and *Irp2*^*-/-*^ mice, which both were elevated (Fig. 5A, B) compared to their sham controls.

**Figure 5:**
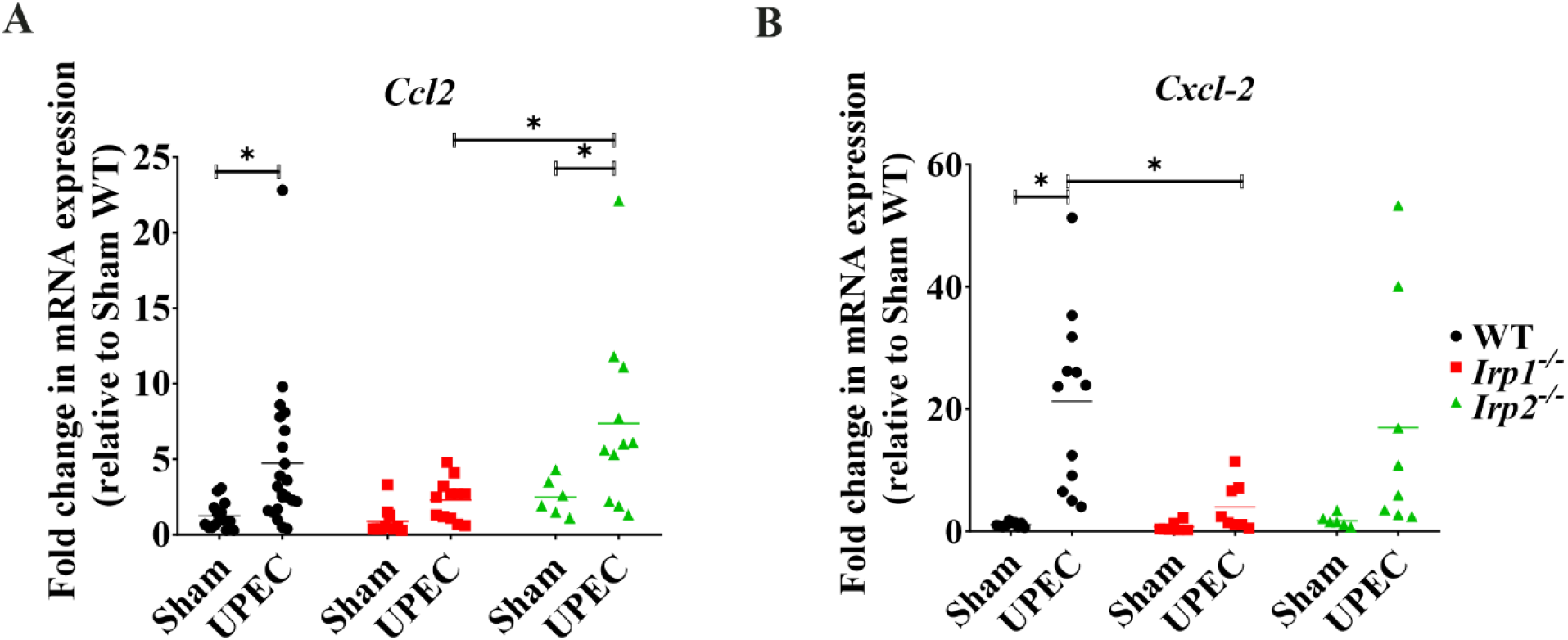
Lower chemokine levels in infected *Irp*1^-/-^ testis compared to WT and *Irp*2^-/-^. **(A, B)** Expression of chemokine levels for the recruitment of monocytes/macrophages (*Ccl2*) and neutrophils (*Cxcl*2) were evaluated by qRT-PCR. Data were obtained from 6-15 mice per group. Statistical significance was determined using two-way ANOVA with Tukey multiple comparison (\**P*<0.05, ** *P*<0.01).

## Discussion

UPEC is a pathogen specialised in survival in extremely iron-poor environments (Dabral et al., 2022; Mann et al., 2017; Mavromatis et al., 2015). The present study provides insights into the role of the iron regulatory proteins IRP1 and IRP2 in modulating the immune response and tissue damage elicited by UPEC-mediated orchitis. In *Irp1-/-* testis the dampened immune response was supported by a decreased ERK1/2 signalling with subsequent lower expression of pro- and anti-inflammatory (*Tnf, Il-*1β, and *Il*-6) cytokines. Based on the critical role of ERK1/2 signalling in immune cell activation and tissue inflammation (Fischer et al., 2005; Lucas et al., 2022), impaired ERK1/2 activation in testis and BMDM from *Irp*1^-/-^ mice suggests a potential regulatory mechanism by which IRP1 interferes with signalling pathways during UPEC infection. This is supported by the notion that ERK interacts with both isoforms of aconitase (ACO1, ACO2) and that inhibition of mitochondrial aconitase interaction with ERK1/2 reduced ERK signalling and the phosphorylation of downstream targets such as p90 ribosomal S6 kinase (RSK) (Talbot et al., 2011).

Previous studies have highlighted the involvement of IRPs in modulating inflammatory responses and immune cell function (Bonadonna et al., 2022; Frost et al., 2022; Nairz et al., 2015). As an example, global disruption of IRPs impairs the development and differentiation of neutrophils in bone marrow of adult mice (Bonadonna et al., 2022). Moreover, IRPs protect the host from *Salmonella* infection by reducing the intracellular proliferation of bacteria through control of iron availability (Nairz et al., 2015). In our study, deletion of IRP2 in naive animals led to a reduction in F4/80^+^CD11b^+^ testicular macrophage numbers without affecting the circulating immune cells. This was accompanied by a massive infiltration of monocytes (Ly6C^+^), macrophages (MHC-II^+^) and neutrophils (Ly6G^+^) in UPEC infected *Irp2*^-/-^ testis, all of which are known for their potential to cause tissue damage through the production of reactive oxygen species (ROS) (Nguyen et al., 2017) and pro-inflammatory cytokines (McBride et al., 2020). We thus suggest, that the noted immune cell recruitment is an instrumental part of the strong inflammatory response and tissue damage observed in *Irp2*^*-/-*^ testis.

In addition to immune cell population dynamics, we investigated the expression of chemokines associated with monocyte (*Ccl2*), macrophage (*Ccl2)* and neutrophil (*Cxcl2)* recruitment. Our qPCR analysis revealed lower expression levels of *Ccl2* and *Cxcl2* in UPEC-infected *Irp*1^-/-^-testis compared highly elevated levels in infected WT and *Irp*2^-/-^-testes (Fig 5). These chemokines play a critical role in the recruitment of immune cells to the site of infection and thus contribute to the inflammatory response. The consistent observation of a lack of significant leukocytic infiltration in the testis of UPEC-infected *Irp1*^-/-^ mice, with the notable exception of the CD206^+^ resident macrophage subpopulation, suggests a role for IRP1 in the recruitment of leukocytes to the testis during infection. These CD206^+^ macrophages are known as immunoregulatory macrophages and play a crucial role in the resolution of tissue inflammation and are established players in the immunoregulation of the testis (Bhushan et al., 2020, 2009; Fijak and Meinhardt, 2006). Their functions extend beyond immune surveillance as they actively contribute to the regulation of local inflammatory processes and the maintenance of tissue homoeostasis. The reduced inflammation and absence of morphological alterations seen in UPEC-infected *Irp*1^-/-^ testis could thus be based on two mechanisms, i.e. the lack of infiltrating proinflammatory cells such as neutrophils and monocytes concomitant with an increase in CD206+ immunoregulatory tissue preserving macrophages. In contrast to the response of *Irp2*^*-/-*^ mice to UPEC-infection, where an exacerbated activation of the ERK pathway was seen in conjunction with higher tissue damage, in *Irp1*^*-/-*^ mice, the immune response seems to be dominated by the reduced ERK signalling.

In summation, our study provides novel insights into the role of IRP1 and IRP2 in modulating the immune response and tissue damage during UPEC-mediated orchitis. These findings suggest the potential of targeting iron regulatory proteins as a therapeutic approach to mitigate testicular damage in bacterial infections. The comprehensive analysis of p-ERK levels, chemokine expression and immune cell dynamics contribute to our understanding of the intricate interplay between iron metabolism, immune signalling and tissue homoeostasis in testicular inflammation. The detailed mechanism remains elusive though why IRP1 and IRP2 that are established to similarly regulate iron homoeostasis have opposing roles in regulating inflammation in UPEC – orchitis.

## Materials and Methods

### Mice

The C57Bl/6J mouse strain was used for all experimental models. *Irp*1^-/-^ and *Irp*2^−/−^ mouse strains were generously provided by Dr. Tracey Rouault (Molecular Medicine Program, National Institute of Child Health and Human Development, National Institutes of Health, Bethesda, MD). Animal experiments were conducted according to ethical approval from the Technion Animal Ethics Committee, Haifa, Israel (IL-135-09-19) and the committee on animal care of the Justus-Liebig-University Giessen (M_819). All animals used in the experiments were age-matched (adult 10 to 12 weeks of age).

### Induction of bacterial orchitis

The uropathogenic *E. coli* (UPEC) strain CFT073 was obtained from ATCC and propagated following established protocols (Michel et al., 2016; Bhushan et al., 2008). To induce an ascending canalicular infection, we performed bilateral ligation of the *vasa deferentia* followed by intravasal injection of UPEC (1×10^5^ colony forming units [CFU] in 10 μl sterile 0.9% NaCl) near the cauda using a Hamilton syringe. The control group (referred to as ‘sham’ mice) underwent the same surgical procedure but received an intravasal injection of 10 μl sterile 0.9% NaCl. Mice were euthanized on day 7 post-infection through isoflurane narcosis followed by cervical dislocation. The selection of this time point was based on previously published report (Michel et al., 2016).

### Histology

The testes were carefully dissected and immediately immersed in Bouin’s fixative (Sigma, cat. no. HT10132-1L, Jerusalem, Israel) for 6 hr. Following fixation, the tissue was embedded in paraffin and cut into 5 μm sections using a microtome and subsequently stained with hematoxylin and eosin.

### Determination of CFU

Testicular samples from both sham and UPEC-infected mice at 7 days post-infection (p.i.) were homogenised in sterile ice-cold PBS (n = 4 to 7 per group). Subsequently, tissue homogenates were subjected to tenfold serial dilutions. Hundred μl of each dilution was streaked onto lysogeny broth (LB) agar plates. The plates were inverted and incubated at 37°C overnight. After incubation, CFUs on each plate were counted, and the values were normalized to tissue weight (per gram of used tissue).

### Immunofluorescence

For immunofluorescence staining of immune cell types in testicular cryosections, 10 μm sections were fixed in methanol at −20°C for 20 min. Following fixation, the sections were washed three times and permeabilised with 0.5% Triton X-100 in 1xTBS for 40 min. Subsequently, blocking was performed with 10% goat serum in 1xTBS for 1 hr. Sections were incubated with an anti-F4/80 antibody (1:100 dilution, Bio-Rad, cat. no. MCA497G, Feldkirchen, Germany or an anti-Ly6G antibody (1:150 dilution, Abcam, cat. no. ab25377, Cambridge, UK) overnight at 4°C. After incubation, the sections were washed three times and incubated with the secondary antibodies (goat anti-rat IgG (H+L) Cross Adsorbed secondary antibody Alexa Fluor546 (Invitrogen, cat. no. A-11081, Karlsruhe, Germany)) for 1 hr at RT. Finally, the sections were mounted, and images analysed on a Zeiss LSM710 Confocal Microscope (Oberkochen, Germany). Adjustments of brightness and contrast on entire images were made using ImageJ. A total of three mice per genotype and treatment were included in the analysis.

### Preparation of single-cell suspension for flow cytometry

Testes were collected into 2 ml tubes containing Dulbecco’s Modified Eagle Medium (DMEM, ThermoFisher, cat. no. 41965039, Qiryat Shemona, Israel) complete media (10% fetal bovine serum (FBS, ThermoFisher, cat. no. 10270106), 1% glutamine (ThermoFisher, cat. no. 25030024), 1% Pen-Strep (ThermoFisher, cat. no. 15140122). For mechanical separation tissues were chopped by scissors. Next, the tissues were digested in DMEM complete media containing 1mg/ml collagenase D (Sigma-Aldrich, cat. no. 11088866001, Jerusalem, Israel) and 1μl DNase (Sigma-Aldrich, cat. no. 4716728001) for 30 min at 37°C under shaking conditions. The cell suspension was further processed by aspiration through a 20G needle and a 70 μm cell strainer (Sigma-Aldrich, cat. No. CLS431751-50EA). The cell suspension was treated with red blood cell (RBC) lysis buffer (Biological Industries, cat. no. 01-888-1B, Beit HaEmek, Israel) to remove RBC. Blood was collected by cardiac puncture and transferred to an anticoagulant blood collection tube (BD, cat. no. BD368841, Elad, Israel). The blood was then subjected to two to three rounds of RBC lysis each for 5 min at RT until no RBCs were visible. The lysis reaction was stopped after each step by adding excess cold PBS, followed by centrifugation at 450xg for 5 min at 4°C. After preparing a single-cell suspension, cells were counted and 1×10^6^ cells/100 μl were used for staining. For live-dead estimation, the cells were first stained with ZombieYellow (15 min at RT, in the dark) and then washed with flow-buffer (0.5 M EDTA pH 8.0; 2.5% bovine serum albumin (BSA) in PBS). The cells were then blocked with anti-mouse CD16/32 for 10 min at 4°C and incubated with the proper antibodies (see Tab. 2 and 3) for 30 min at 4°C.

After the incubation, cells were washed with flow-buffer and fixed with 1% paraformaldehyde (PFA, Barnaor, cat. No. BN15710, Petah Tikva, Israel) for 15 min at 4°C. After washing with cold PBS, the cells were resuspended in flow-buffer for analysis on the Cytec® Aurora Spectral Flow cytometer, which was kindly made accessible by the Technion Life Sciences and Engineering Infrastructure Centre (LS&E) in Haifa, Israel. The data were analysed with FlowJo software version 10.8.1.

**Table 1:** Macrophage and neutrophil panel.

| Antibodies and viability dye | Clone | Fluorophore | Supplier |
| --- | --- | --- | --- |
| MHC-II (I-A/I-E) | M5/114.15.2 | PE-Cy7 | Biolegend; Cat. No. 107629 |
| CD11b | M1/70 | APC-Cy7 | Biolegend; Cat. No. 101226 |
| Ly6-g | 1A8 | BV421 | Biolegend; Cat. No. 127627 |
| Ly6-c | HK1.4 | APC | Biolegend; Cat. No. 128015 |
| CD45 | I3/2.3 | FITC | Biolegend; Cat. No. 147710 |
| CD206 | C068C2 | PE | Biolegend; Cat. No. 141705 |
| F4/80 | BM8 | PE-Cy5 | Biolegend; Cat. No. 123111 |
| CD11c | N418 | BV510 | Biolegend; Cat. No. 117337 |
| ZombieYellow |  |  | Biolegend; Cat. No. 423105 |
| CD16/32 | Clone 93 | - | Biolegend; Cat. No. 101302 |

**Table 2:** B and T cell panel.

| Antibody and viability dye | Clone | Fluorophore | Supplier |
| --- | --- | --- | --- |
| CD19 | 6D5 | Pacific Blue | Biolegend; Cat. No. 115523 |
| CD11b | M1/70 | APC-Cy7 | Biolegend; Cat. No. 101226 |
| CD3 | 17A2 | APC | Biolegend; Cat. No. 100235 |
| CD45 | 30-F11 | PE | eBioscience; Cat. No. 12-0451-82 |
| ZombieYellow |  |  | Biolegend; Cat. No. 423105 |

### SDS-PAGE and Western Blot

Tissue samples were immediately collected and snap-frozen at −80°C for subsequent analysis. Approximately 10mg of testis tissue was homogenized in RIPA-Buffer (50 mM Tris-HCL pH 7.4, 150 mM NaCl, 0.5% deoxycholate, 0.1% SDS, 1% NP-40, AEBSF 0.1 M, 1 mM DTT) supplemented with phosphatase inhibitor (Roche, cat. No. 4906845001, Basal, Switzerland) and protease inhibitor cocktail (Roche, cat. No. 11836170001) followed by incubation on ice for 30 min. After incubation, the samples were centrifuged at 20,000xg for 20 min at 4°C, and the supernatant was collected in a new tube. The protein concentration in the supernatant was measured by the BCA protein assay kit (Sigma-Aldrich, cat. No. BCA1-1KT).

Equal amounts of protein (20-50 μg) were separated by 10-15% SDS-PAGE. Following electrophoresis, the proteins were transferred onto PVDF membrane (Merck Millipore, cat. no. IPVH00010, Jerusalem, Israel). Prior to antibody incubation, the membranes were blocked with blocking solution (Tris-buffered saline with 0.1% Tween-20 supplemented with milk) for 1 hr at RT to reduce non-specific binding of antibodies. For primary antibody incubation, the appropriate antibodies (see Table 5) were diluted in antibody buffer (2.5% BSA in TBST and 0.02% sodium azide) and the membranes were incubated overnight at 4°C. After washing with TBST buffer, the membranes were incubated with horseradish peroxidase-conjugated α-rabbit (Abcam, cat. no. ab97200, Cambridge, UK) or polyvalent α-mouse (DanyelBiotech, cat.no. NXA931, Rehovot, Israel) immunoglobulin secondary antibody for 1 hr at RT. Antibody detection was performed using the ECL Plus chemiluminescence Western Blot kit (Advansta, cat. no. K-12042-D20, San Jose, USA).

### Quantification of mRNA by qPCR

RNA was isolated from snap-frozen testes using Trizol (Invitrogen, cat. no. 15596-018, Karlsruhe, Germany) following the manufacturer’s protocol. To eliminate genomic DNA contamination, a DNase digestion step was performed using the PerfeCTa Dnase I kit (Quanta Bioscience, cat. no. 95150, Yakum, Israel) according to the manufacturer’s instructions. Subsequently, 1 μg of RNA was reverse transcribed to cDNA using the qScript cDNA synthesis kit (Quanta Bioscience cat. no. 95048) as per the manufacturer’s protocol. Quantitative PCR was carried out on a QuantStudio 12K Flex machine (Applied Biosystem, Rhenium, Israel) using PerfeCTa SYBRGreen SuperMix (Quantabio, cat. no. 95074-012-4,). The primers utilised in this study are listed in Table 5. The relative accumulation level of each mRNA was normalised to *RplpO* (for tissue) as a reference, and the comparative Ct method (ΔΔCt method) was employed for the data analysis.

### DNA extraction from FFPE-embedded tissue

DNA was extracted from paraffin-embedded testes to quantify total bacterial load 7d p.i.. A total of eight sections (5 μm thick) were collected in a 2ml tube. DNA extraction was performed using the QIAamp DNA FFPE Tissue Kit (Qiagen, Cat. no. 56404, Hilden, Germany) following the manufacturer’s instructions. As demonstrated in previous studies (Klein et al., 2020, 2019; Lu et al., 2013; Michel et al., 2016) the isolated DNA (N=5) was used for qPCR analysis targeting the bacterial marker gene PapC (primer sequence in Table 5). The relative abundance of each mRNA was normalised to *actin* as a reference, and the comparative Ct method (ΔΔCt method) was employed for data analysis.

### Isolation of bone marrow cells

Bone marrow-derived macrophages (BMDM) were generated following the protocol described by (Haag and Murthy, 2021)with slight modifications. Briefly, bone marrow cells were obtained by flushing the femurs and tibias from C57BL/6 mice (WT, *Irp1*^*-/-*^ and *Irp2*^*-/-*^). After RBC lysis, the cells were plated in DMEM medium supplemented with 20% FCS (Biological Industries), 30% CCL1 cell-conditioned medium (L929 cells were a kind gift from Dr. J. Kaplan, University of Utah) 1% L-glutamine and 1% Pen-Strep. The cells were incubated for 6 days at 37°C, 5 % CO_2_ and 6% O_2_. On day 6, fully differentiated macrophages were harvested. On experimental day cells were stimulated with 200 ng/ml lipopolysaccharide (LPS, Sigma-Aldrich, cat. no. L4391) or without LPS in a time dependent fashion. Extracted proteins from BMDM were for Western blot analysis.

### Statistics

Statistical analyses were performed using the GraphPad Prism software (version 8.0). The data from infected samples were analysed using two-way analysis of variance (ANOVA) followed by Tukey’s multiple comparisons was used. The non-infected samples were statistically analysed using the Kruskal-Wallis test followed by Dunn’s multiple comparisons. A probability value of *P* < 0.05 was considered statistically significant. Statistical significance levels were denoted as \**P*<0.05, ** *P*<0.01.

## Acknowledgements

We would like to express our gratitude to Dr. T. A. Rouault (NICHD, NIH, Bethesda MD, USA) for generously providing the *Irp*1^-/-^ and *Irp*2^-/-^ mice used in this study. Additionally, we would like to extend our thanks to the Technion Life Sciences and Engineering Infrastructure Centre (LS&E) in Haifa, Israel for granting us access to the Cytec® Aurora Spectral Flow cytometer and for valuable technical support.

## Author contributions

NG and AH contributed to the performance, analysis, validation, and supervision of the *in vivo* experiments as well as writing the manuscript. BA performed the BMDM experiments. SB and NGR provided intellectual support to the concept and contributed to the editing and review of the manuscript. EGMH and AM were involved in the conceptualization of the study, supervision, acquisition of funding and contributed to the writing and editing of the manuscript.

---

**Table 3:** Antibodies used for Western Blot.

| Antibodies | Source | Supplier |
| --- | --- | --- |
| Ferritin-L | Rabbit | Abcam, Ab69090, Cambridge, UK |
| Ferritin-H | Rabbit | Cell signaling, 4393S, Cambridge, UK |
| p-RELA/NFκB p65 Antibody (27.Ser 536) | Mouse | Santa Cruz, SC-136548, Petach Tikvah Israel |
| NF-κB p65 (D14E12) (Ser536) | Rabbit | Cell signaling, 8242, Cambridge, UK |
| Phospho-p44/42 MAPK (Erk1/2)<br>(Thr202/Tyr204) | Rabbit | Cell signaling, 9101S, Cambridge, UK |
| p44/42 MAPK (Erk1/2) | Rabbit | Cell signaling, 9102S, Cambridge, UK |
| Phospho-p38 MAPK (Thr180/Tyr182) | Rabbit | Cell signaling, 4511, Cambridge, UK |
| p38 MAPK | Rabbit | Cell signaling, 8690, Cambridge, UK |
| alpha Tubulin | Mouse | Santa Cruz, SC-69969, Petach Tikvah Israel |

**Table 4:**
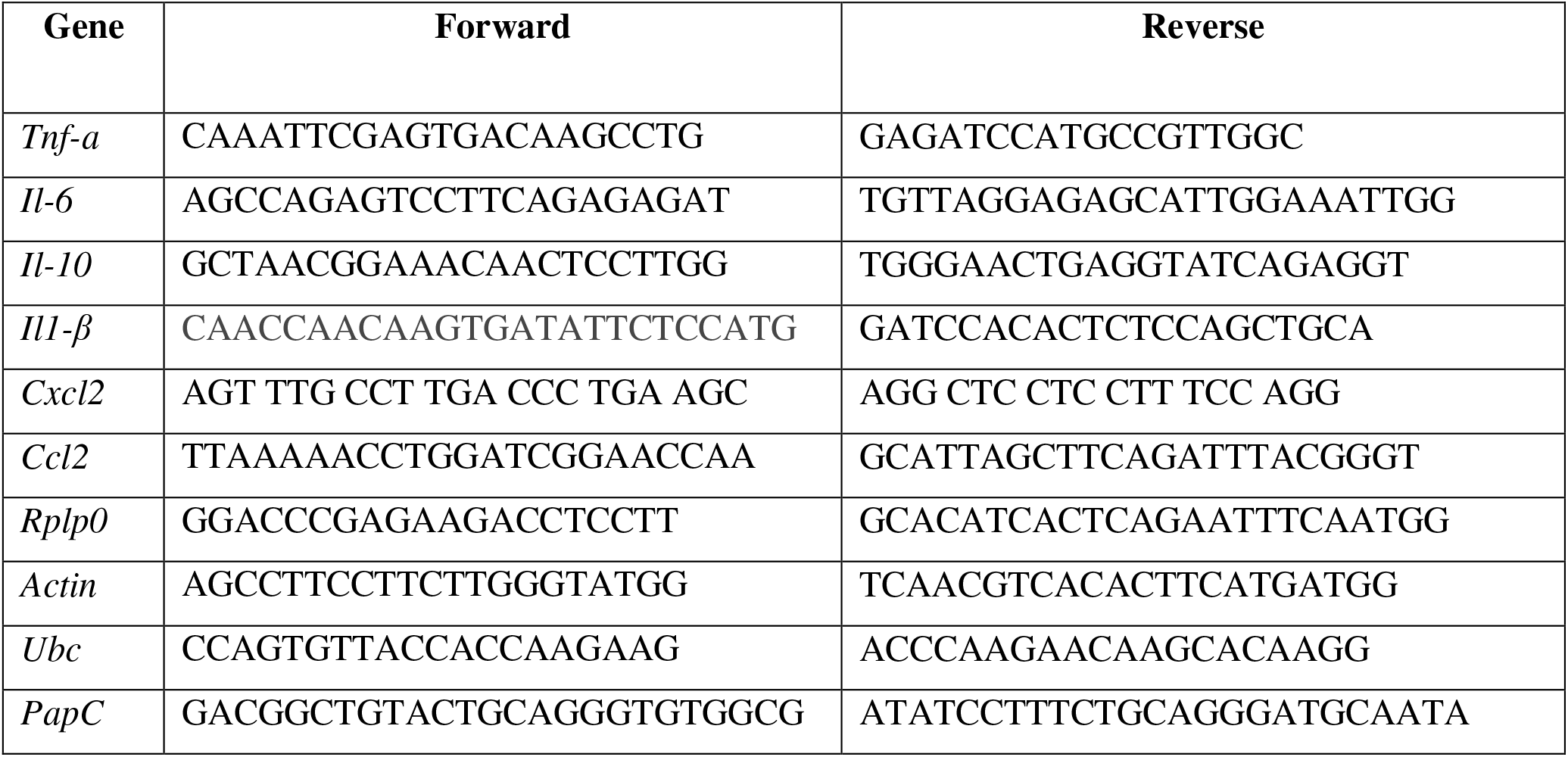
Sequences of primers used for qPCR.

