## Supplementary data for "Iron regulatory proteins 1 and 2 have opposing roles in regulating inflammation in bacterial orchitis"

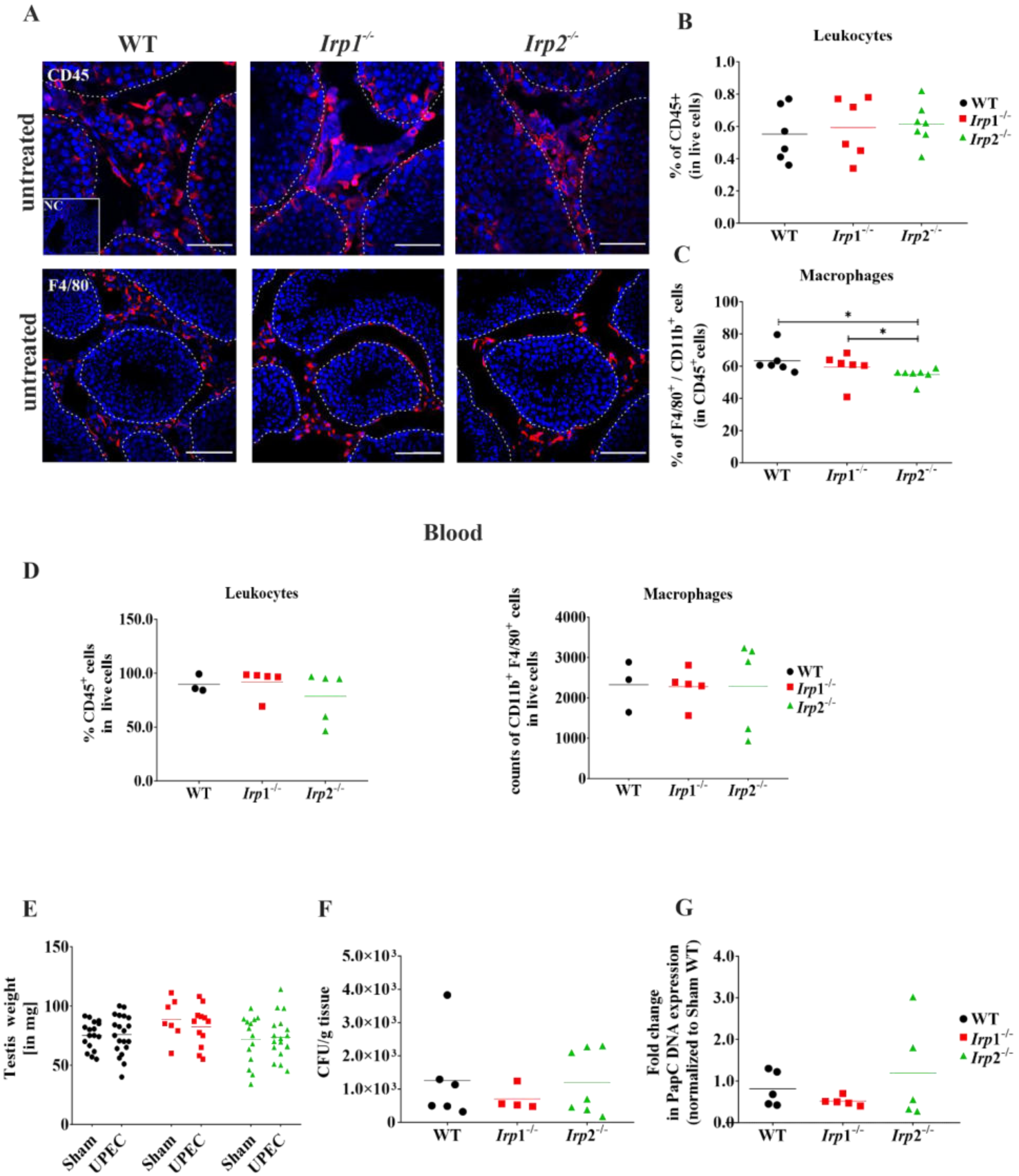


**Supplementary Figure S1** (A) Immunofluorescence staining of cryosections and (B, C) flow cytometry analysis of untreated testis samples of all three genotypes (WT, *Irp1*$^{-/-}$, *Irp2*$^{-/-}$) were conducted using CD45 and CD11b and F4/80 markers to identify changes in total leukocytes and macrophages. Representative immunofluorescence images of total leukocytes (CD45) and macrophages (F4/80) are provided (N=3-6, scale bar=100µm). Statistical analysis was performed using the Kruskal-Wallis test followed by Dunn´s multiple comparisons. (D) Analysis of circulating immune cells was performed using flow cytometry on blood samples collected from WT, *Irp1*$^{-/-}$ and *Irp2*$^{-/-}$ mice. Representative graphs of total leukocytes ($CD45^+$; in $2x10^5$ live cells) and macrophages ($F4/80^+$ $CD11b^+$; in $2x10^5$ live cells) are shown. (F) Testis weights were recorded after 7 days of infection, (F; G) and the bacterial load in testis were quantified using (F) CFU and (G) PapC qPCR analysis. A summary of three independent experiments is provided, with N= 4–7 for CFUs and N=5 for PapC qPCR per group.

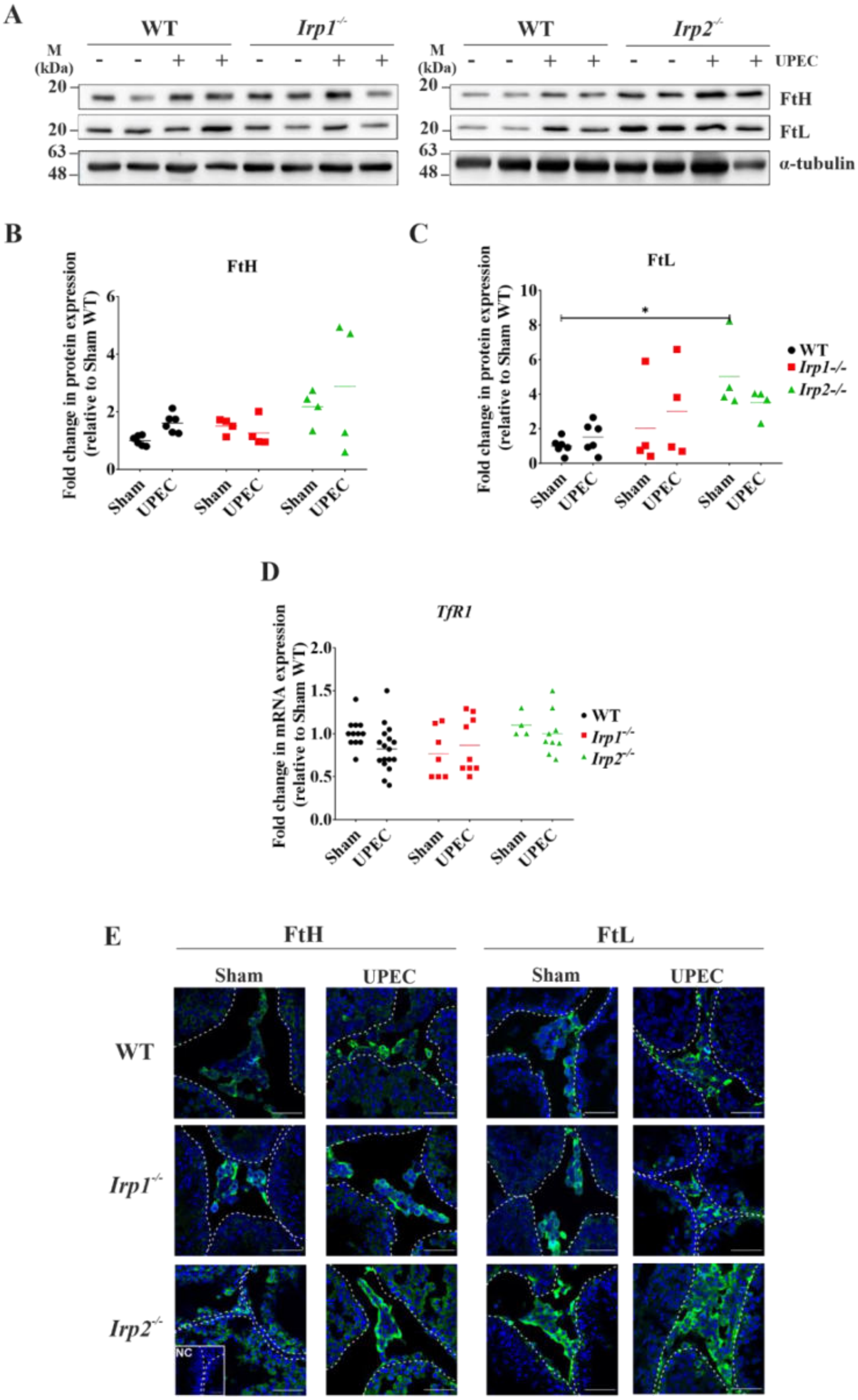


**Supplementary Figure S2.** (A, B) Protein levels of ferritin-H (FtH) and ferritin-L (FtL) in UPEC-infected WT, *Irp1*$^{-/-}$ and *Irp2*$^{-/-}$ testis were analysed by Western blot. The band intensities of FtH

(B) and FtL (C) were quantified using ImageJ and normalised to $\alpha$-tubulin. Representative images are shown. Significant differences are indicated (*$P<0.05$, ** $P<0.01$), N=4-6 independent samples. (D) Quantitative RT-PCR analysis of transferrin receptor (*Tfr1*) in UPEC-infected testis (N= 6-15 mice per group). (E) Immunofluorescence staining was performed to investigate the changes in the localisation of FtH and FtL in UPEC-infected testes. Cryosections of all three genotypes were stained for FtH (green), FtL (green) and DAPI (blue). Representative images are shown. Scale bar = 50 μm, N=5.

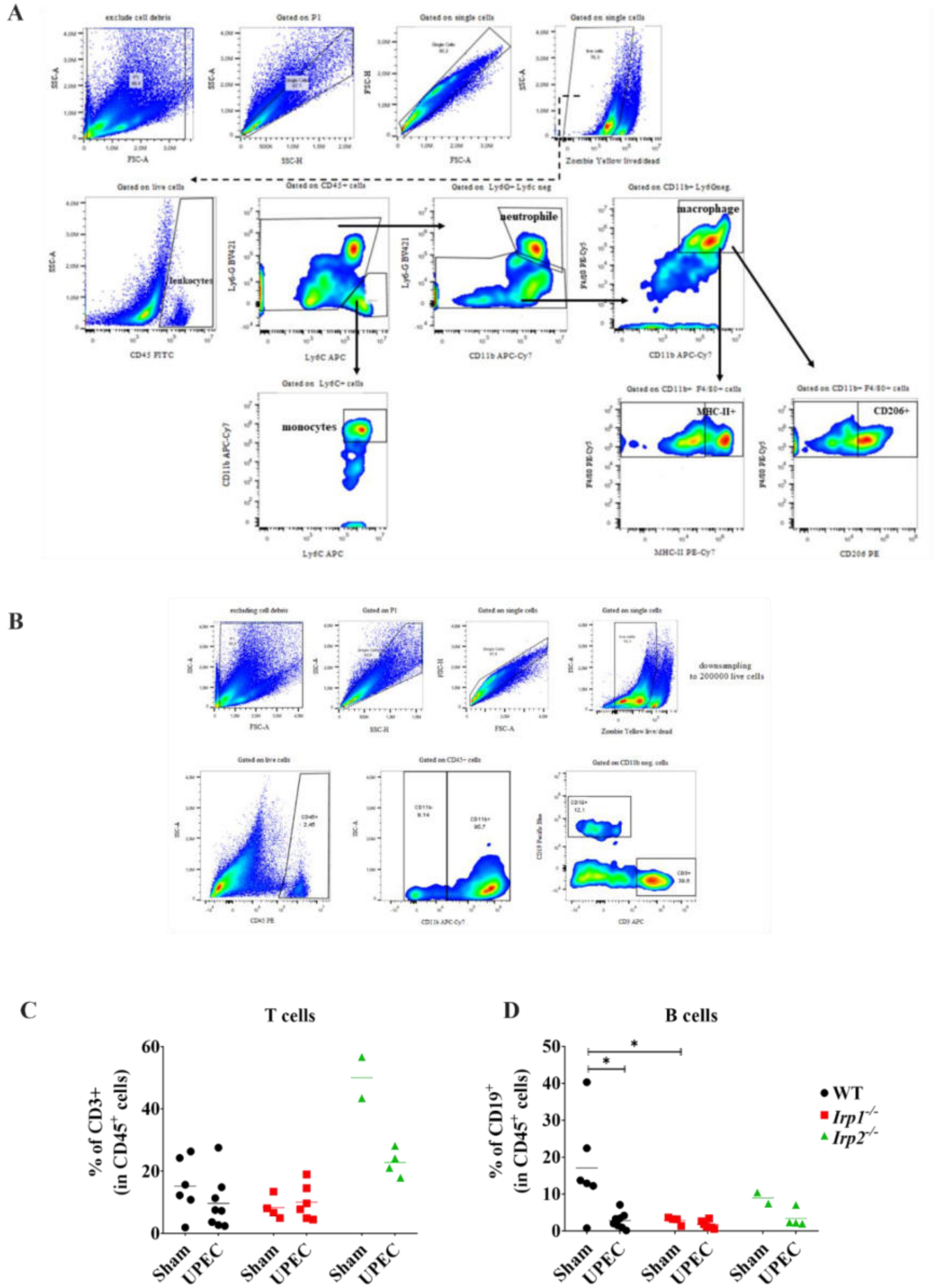


**Supplementary Figure S3:** (A, B) Gating strategy for the macrophage and neutrophil antibody panel in infected testis samples. Live cells were initially gated for CD45 to select total leukocytes.

From the $CD45^+$ population, monocytes were further gated using Ly6G and Ly6C expression. The gate of $Ly6G^+$ and Ly6C- cells was then used to identify granulocytes using CD11b and Ly6G expression. After excluding the monocytes and granulocytes, macrophages were gated ($CD11b^+Ly6G^-$). Finally, macrophages and their subpopulations were identified using the following markers.: $CD11b^+F4/80^+$, $F4/80^+MHC\text{-}II^+$, and $F4/80^+CD206^+$. (B-D) Gating strategy and respective graphs of B and T cells are presented. Live cells were initially gated for CD45 followed by gating on CD11b alone (B) with subsequent selection of the $CD11b^-$ population for gating on CD19 (B cells) and CD3 (T cells) expression, respectively. The graphs depict the percentages of B cells (C) and T cells (D) within the $CD45^+$ population. Data were obtained from infected testis samples and represent the summary of three independent experiments (each N= 2–3 per group). Statistical significance was determined using two-way ANOVA with Tukey multiple comparison (*$P$<0.05, ** $P$<0.01).

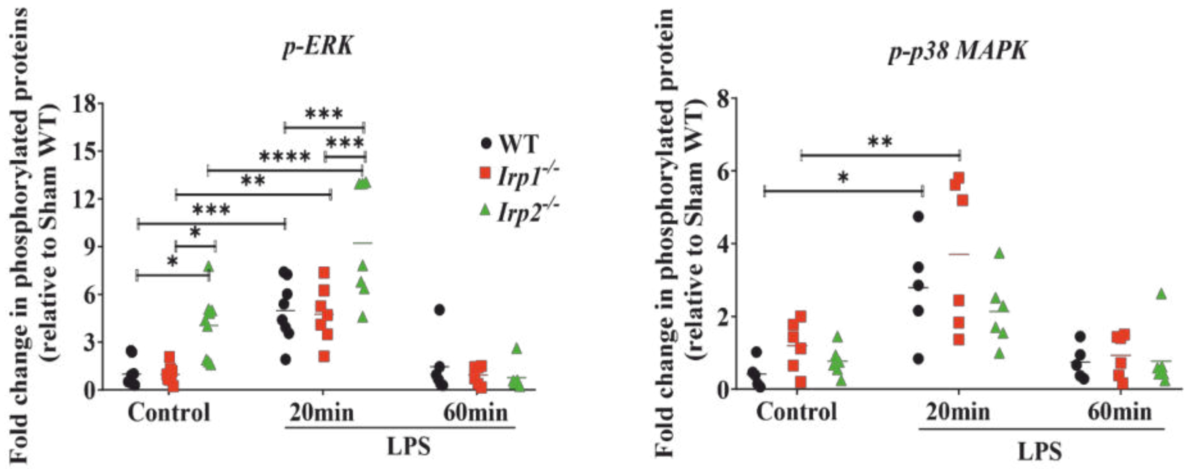


**Supplementary Figure S4:** Bone marrow-derived macrophages (BMDM) from WT, *Irp1*$^{-/-}$ and *Irp2*$^{-/-}$ mice were isolated and stimulated with 200 ng/ml lipopolysaccharide (LPS) for different time points (20 and 60 min) under specific oxygen conditions (6% $O_2$ at 37°C). Protein levels of indicated signalling molecules were analysed by Western blot. Representative images are shown. Quantitative analysis of the band intensities of p-ERK and p-P38 were performed using ImageJ and normalised to the corresponding loading control. Statistical analysis revealed significant differences (*P<0.05) between the genotypes and time points.
